# *decepentaplegic* directs wiring of female-differentiated *fruitless* sex peptide response-inducing neurons

**DOI:** 10.64898/2026.05.06.723318

**Authors:** Deepanshu N.D. Singh, Matthias Soller

## Abstract

Female reproductive success requires coordinated behavioural responses following mating. In *Drosophila melanogaster*, these responses are induced by male-derived sex-peptide (SP) and include reduced receptivity to further mating and increased oviposition. Although the neural pathways controlling these behaviours have been partially characterized, the developmental mechanisms that establish and maintain these circuits remain poorly understood. Using a genetic approach, we identified an EMS-induced mutant that retains eggs and fails to reduce receptivity following SP exposure. We mapped this mutation to the transcription termination zone after the polyA site of the *dpp* locus and show that this allele *dpp^HB3^* affects expression. Dpp/BMP signalling acts within SP response-inducing neurons (SPRINz) and is required for correct neuronal wiring in *dpp^HB3^* mutants. We further find that the SPRINZ-*fru11/12* enhancer induces gene expression upon Dpp exposure. Also, female sexual differentiation trough the sex determination gene *tra* is required in SPRINz to display post-mating behaviours. Together, these findings suggest a new role for *dpp* in specifying neuronal connectivity in the context of sexual differentiation by the *Drosophila* canonical sex determination pathway to implement neuronal wiring for post-mating behaviours.

## Introduction

Male-derived factors transferred during mating regulate female physiology and behaviour in most insects (Gillott, 2003; Avila et al., 2011; Singh and Soller, 2025). In *Drosophila*, mating triggers a characteristic set of female-specific responses that include reduced receptivity to further mating and increased egg laying (Kubli, 2003). A key signal driving these post-mating responses is male-derived sex peptide (SP). SP is a 36 amino acid peptide that is transferred to females during copulation.

SP induces broad behavioural and physiological changes beyond reducing receptivity and increasing oviposition. These changes include enhanced egg production, elevated feeding, altered food preference, modulation of sleep and memory, constipation, activation of the immune system and impacts on life span as cost of mating (Soller et al., 1999; Peng et al., 2005a; Carvalho et al., 2006; Domanitskaya et al., 2007; Isaac et al., 2010; Ribeiro and Dickson, 2010; Cognigni et al., 2011; Scheunemann et al., 2019). SP also binds to sperm and functions as a molecular sensor for sperm storage. This interaction is required for the release of stored sperm (Peng et al., 2005b; Wigby and Chapman, 2005; Avila et al., 2010). Moreover, SP binding to sperm extends the SP response from one to about seven days (Chapman et al., 2003; Liu and Kubli, 2003).

The downstream effects of SP on female physiology are well documented, yet key features of the neuronal circuitry that regulate core post-mating behaviours remain incompletely understood. Reduced receptivity and increased oviposition represent the central behavioural outputs of the post-mating response. Availability of the *Drosophila* neuro-connectome will in principle allow to identify the exact neuronal connections linking receptor activation to motor programs (Scheffer et al., 2020; Phelps et al., 2021; Galili et al., 2022; Nallasivan et al., 2026).

Initial insights into the neuronal circuitry underlying the sex peptide response came from egg retainer mutants insensitive to sex peptide (Soller et al., 2006). Mutations in the *egghead* gene, which encodes a 1,4-mannosyltransferase required for glycosphingolipid biosynthesis, display defects in neuronal wiring in connections of the ventral nerve cord to the central brain (Fan et al., 2005; Soller et al., 2006).

A key tool to map target neurons for neuropeptides is based on targeted expression of membrane-tethered peptide (Nakayama et al., 1997; Choi et al., 2009; Choi and Nitabach, 2013). Although initial screens for SP target neurons using enhancer-trap *GAL4* lines identified only broad expressing lines (Nakayama et al., 1997), neurons expressing the sex determination genes *fruitless (fru),* and *doublesex (dsx)* can induce the SP post-mating response from expression of membrane-tethered SP (mSP) (Kvitsiani and Dickson, 2006; Yapici et al., 2008; Rezaval et al., 2012). Intriguingly, even-though both reduction of receptivity and induction of egg laying are allosterically induced by the same critical concentration of SP, these two responses can be separated (Haussmann et al., 2013).

Recent development of tools for intersecting gene expression patterns has substantially increased the resolution for expression of mSP or RNAi of Sex peptide receptor (SPR) in only few neurons (Nallasivan et al., 2026). Such approach using systematic tiled regulatory regions in *SPR*, *fru* and *dsx* genes identified several distinct populations of SP response–inducing neurons *(SPRINz)* in the central brain and abdominal ganglion (Nallasivan et al., 2026). However, how these circuits are established during development and how they are linked to the sex determination pathway remains unknown.

Here, we identified a hypomorphic allele (*HB3*) of *decapentaplegic* (*dpp*) in a genetic screen for sex peptide insensitive egg-retainer mutants (Soller, 2006; Schupbach, 1991). Dpp as a member of the Bone Morphogenetic Protein (BMP) pathway is a conserved regulator of growth, differentiation, and morphogenesis across animals (Affolter and Basler, 2007; Hamaratoglu et al., 2014; Matsuda et al., 2021; Matsuda and Affolter, 2023). In *Drosophila*, Dpp acts as a morphogen to provide positional information within developing tissues and directs cell proliferation, pattern formation, and spatial organization (Affolter and Basler, 2007) (Nellen et al., 1996; Vuilleumier et al., 2010; Peterson et al., 2022). Dpp binds to a heteromeric receptor of *thickveins* and variable partners including *wishfull thinking*, that upon activation by Dpp regulate transcription through the transcription factor *mothers against Dpp* (Yamashita et al., 1996). However, the role of *dpp* in nervous system development remains largely unexplored. Here, *dpp* has roles in the specification and development of the *pars intercerebralis* and *pars lateralis*, neuroendocrine command centres in the *Drosophila* brain (de Velasco et al., 2007), connectivity of central pacemaker neurons (Polcowñuk et al., 2021) and in the control of synapse numbers (Marqués et al., 2002; Haussmann et al., 2008).

Molecular genetic analyses reveal that the *dpp^HB3^* lesion after the polyA site impacts on *dpp* expression and on neuronal wiring in *fru11/12* SPRINz. Moreover, masculinizing SPRINz reveals that specification is under the control of the sex determination pathway to implement the female sex peptide induced post-mating response.

## Results

### *dpp^HB3^* mutant females retain eggs and are insensitive to sex-peptide

Egg retention and remating are core behavioural readouts of SP insensitivity (Soller et al., 2006). To identify novel genes specifying SP-mediated responses, we screened females from egg retainer lines. These females exhibit normal oogenesis yet fail to lay eggs and do not appropriately reduce receptivity following mating (Soller et al., 2006). Using this approach, a homozygous viable EMS-induced mutant line, *HB3* (Schupbach and Wieschaus, 1991), was identified that retained eggs and failed to reduce receptivity after SP injection (Fig. 1 B,C). We mapped this allele by meiotic recombination based on the egg retention phenotype measured by ovulation relative to cinnabar and brown markers to 2-41.8, but the location is further away from the centromere because egg retention is not complete (Nallasivan et al., 2021). To identify the locus for this EMS allele, we then crossed the Bloomington deficiency kit for the left arm of chromosome 2 to *HB3* and identified two deficiencies which displayed a phenotype when transheterozygous with *HB3*. *Df(2L)Exel7011* resulted in lethality (n=345 flies analysed) (Fig. 1A) and *Df(2L)ED4651* resulted in sterility (n=56 sterile females) (Fig. S1A), but we did not find the anticipated egg retention phenotype for this second locus. Using overlapping deficiencies and null-mutant alleles lethality could then be mapped to the *decapentaplegic* (*dpp*) locus in chromosome section 22F1-3 (2-6, *dpp^hr56^*, n=213). To demonstrate that *HB3* is indeed an allele of *dpp*, we combined a *UASdpp* into *HB3*/*dpp^hr56^*. The leaky expression of the *UASdpp* insert rescued lethality of *HB3*/*dpp^hr56^*(83%, n=42) and these females laid eggs and were fertile (Fig. 1D,E). Moreover, these females responded normally to SP by increasing egg laying and reducing receptivity and we now refer to this allele as *dpp^HB3^* (Fig. 1D,E).

**Fig. 1:**
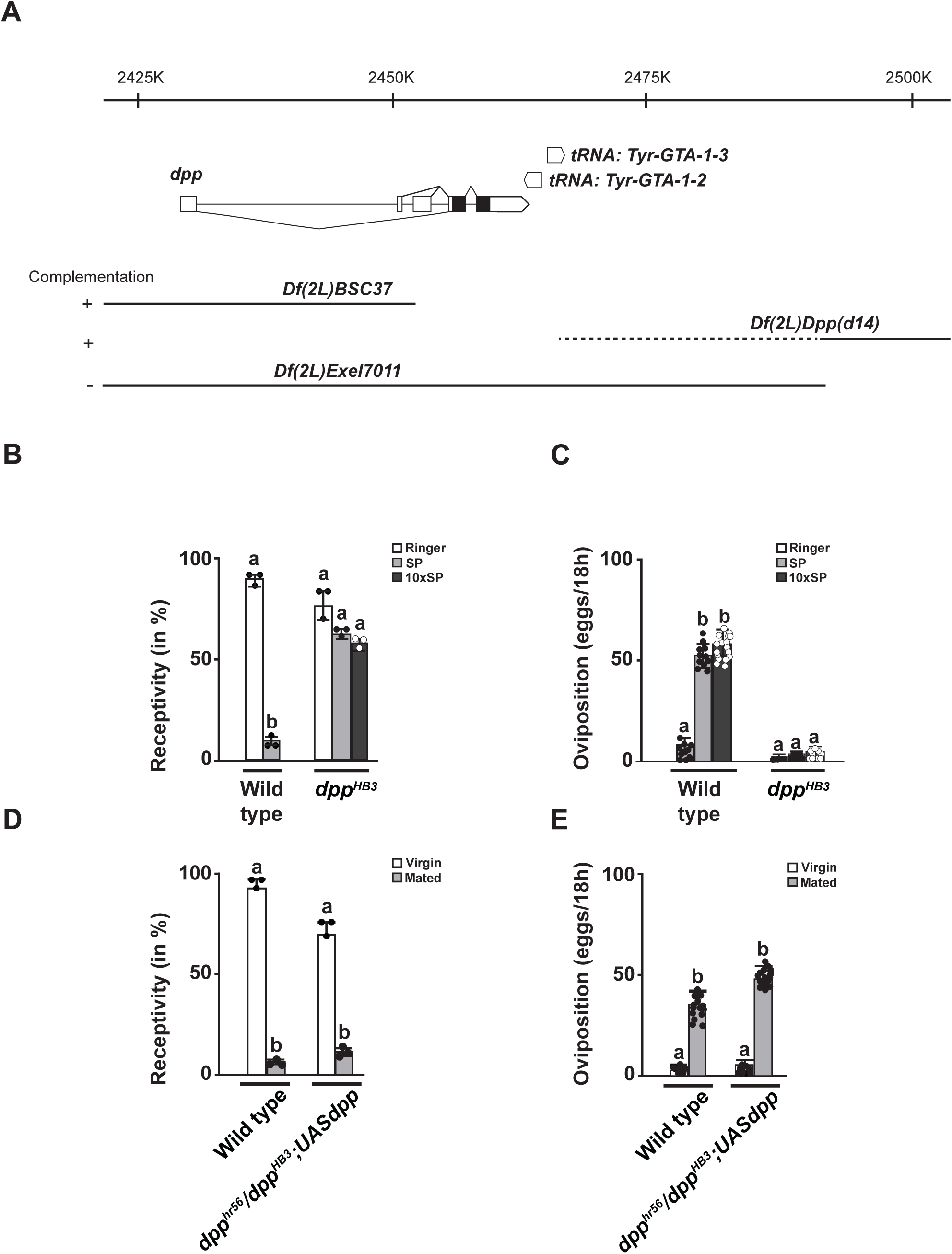
Mapping of the sex-peptide insensitive EMS allele *HB3* to *dpp*. A) Schematic of the *dpp* chromosomal region depicting gene models and chromosomal deficiencies depicted below chromosomal nucleotide positions. Coding parts are shown as black and non-coding parts as white boxes. Complementation data is shown on the left. B and C) Receptivity and oviposition of wild type, homozygous *dpp^HB3^* mutants after sex-peptide (SP, black) or Ringer’s (R, white) injection measured by counting mating females in a 1 h time period 3 h after SP or R injection. D and E) Receptivity and oviposition of wild type transheterozygous females *dpp^hr56^/dpp^HB3^*; *UASdpp/+*, (Virgin, white) and (mated, black), respectively for receptivity, or for oviposition, means of eggs laid in 18 h with the standard error for 10 females each. For receptivity means with the standard error for three or four experiments with 18-23 females each are shown. Statistically significant differences from ANOVA post-hoc pairwise comparisons are indicated by different letters (p≤0.0001).

The sterility locus mapped to chromosome section 23F3 uncovered by *Df(2L)drm-P2* and *Df(2L)Exel7016*, but we were not able to map the lesion further using transposon inserts in *ND-PDSW* (lethal *P{lacW}ND-PDSW^k10101^*), *Pif1 (P{EPgy2}Pif1^EY10295^*) and *Pgant4* (lethal *Mi{ET1}Pgant4^MB04930^*) (Fig. S1A).

To identify the molecular lesions in *dpp^HB3^* in these two genomic regions, we prepared DNA from mutant *dpp^HB3^* flies and sequenced the whole genome and focused on lesions that are not annotated as polymorphisms in the Drosophila Genetic Reference Panel (DGRP2) (Mackay et al., 2012; Huang et al., 2014). In both *dpp* and the *msl-2* loci, we did not find changes in the protein coding region. In the *dpp* locus, we identified deletions of 9 nt and 3 nucleotides distal of the polyA site (GGTGAGGGATA**<u>AAAGAAAAG</u>**TATA**<u>TGG</u>**TATACA) (Fig. 2A). In the 23F3 region, we identified multiple changes in *msl-2* near the Sxl binding sites in the 3’UTR (TTTTTTTGA**<u>A</u>**GCA, TTTTTTTTTAA**<u>AAA</u>**TAGG and TTTTTTTTTAA**<u>A</u>**TACG, A insertions are underlined) and in the polyA site (AAATAA**<u>A/C</u>**A), but females transheterozygos for the *msl-2^227^*null allele and *dpp^HB3^* were not sterile and did not retain eggs (Fig. S1A).

**Fig. 2:**
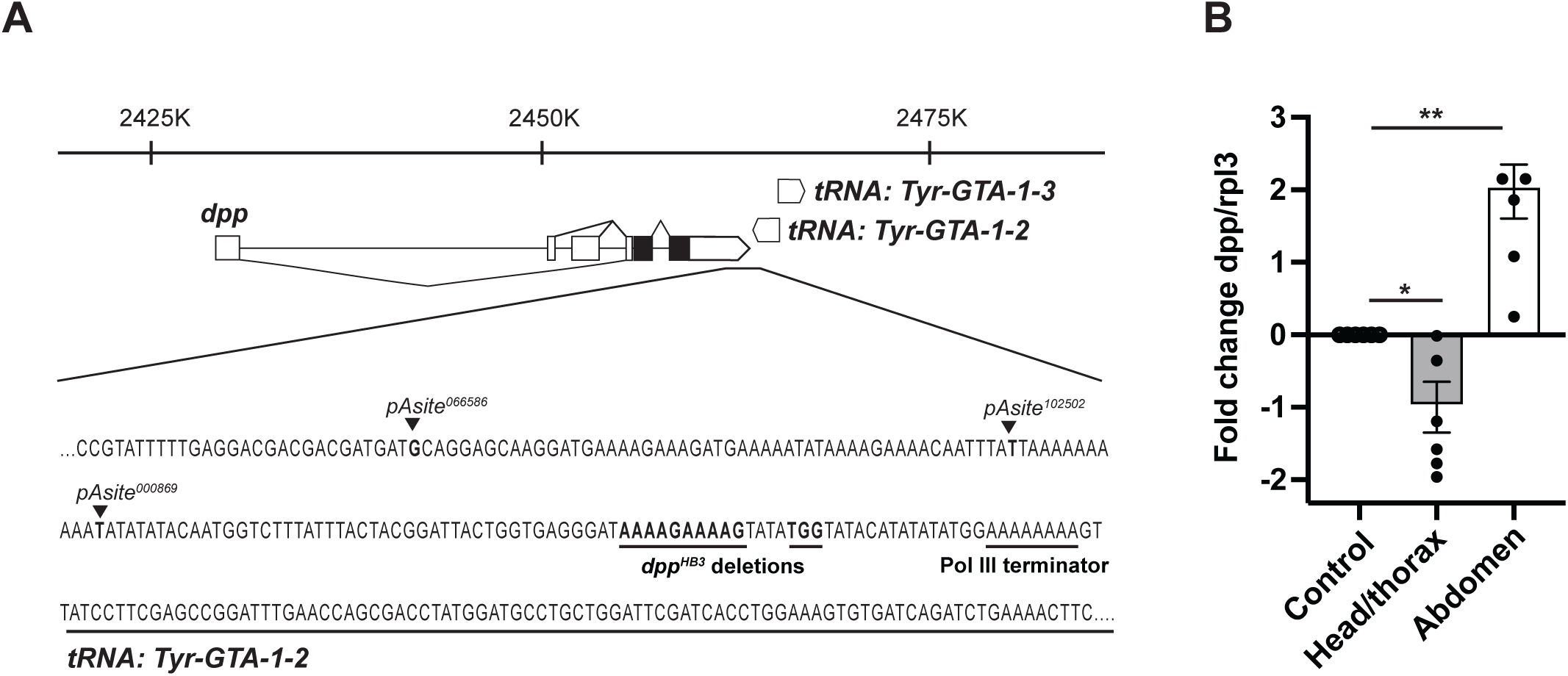
*dpp* expression is reduced in *dpp^HB3^* females. **A)** Schematic of the *dpp* chromosomal region depicting gene models with coding parts are shown as black and non-coding parts as white boxes. The sequence spaning the *dpp* 3’ part into the adjacent tRNA gene is shown at the bottom. Annotated polyA sites (polyA addition) are from Flybase. Nucleotide deletions in *dpp^HB3^*are shown in bold and underlined. The tRNA gene is underlined and the RNA Polymerase III terminator sequence indicated. B) Expression of *dpp* polyA mRNA determined by qRT-PCR is shown as fold change normalized to *rpl3* as a housekeeping control in head/thorax (grey bar) and abdomen part of the fly (white bar). DeltaCT mean ± standard deviation. Significance from unpaired student t test for the head/thorax part is *p < 0.0488 and for the abdomen part is **p < 0.0082 (from 3 biological replicates each).

### *dpp* expression is reduced in *dpp^HB3^* mutant females

The molecular lesion in *dpp^HB3^* maps distal to the polyA site within the binding region of the core polyadenylation factor CstF64 (Soller and White, 2003; Soller, 2006). Because *dpp^HB3^* behaves as a hypomorphic allele, we tested whether *dpp* expression is reduced in *dpp^HB3^*mutants using quantitative RT–PCR. Indeed, *dpp* transcript levels were reduced in the head and thorax of HB3 mutants (Fig. 2B). In contrast, *dpp* expression was increased in the abdomen.

### RNAi knockdown of key Dpp/BMP signalling pathway components compromises the post-mating response in *fru11/12* and *FD6* sex peptide response-inducing neurons

Expression of mSP by intersection of expression patterns using splitGAL4 identified five populations of sex peptide response inducing neurons (SPRINz, *SPR8 ∩ dsx*, *SPR8 ∩ fru11/12*, *SPR8 ∩ FD6*, *fru11/12 ∩ dsx* and *fru11/12 ∩ FD6*) (Nallasivan et al., 2026). To evaluate whether Dpp/BMP signalling directly affects the sex peptide response in SPRINz, we downregulated key components of the Dpp/BMP signalling pathway using RNAi. For each experiment, we chose the strongest RNAi line based on wing development defects from expression with *enGAL4*, but with no other adverse effects. BMP receptors exist in two forms, type I and type II, both single-pass transmembrane serine/threonine kinase receptors that assemble into a heterotetrameric signalling complex composed of two type I and two type II receptors (Yamashita et al., 1996). Activation of the receptor by Dpp induces a signalling pathway resulting in transcriptional changes induced by the transcription factor *mothers against dpp* (*mad*).

We first downregulated the type I receptor *thickveins* (*tkv*) using *tkvRNAi* in the SPRINz *split-GAL4* lines. Among the five lines tested, only *SPR8 ∩ FD6* and *fru11/12 ∩ dsx* showed a significant increase in remating (Fig. 3A), whereas oviposition was significantly reduced across all five lines relative to mated controls (Fig. 3B).

**Fig. 3:**
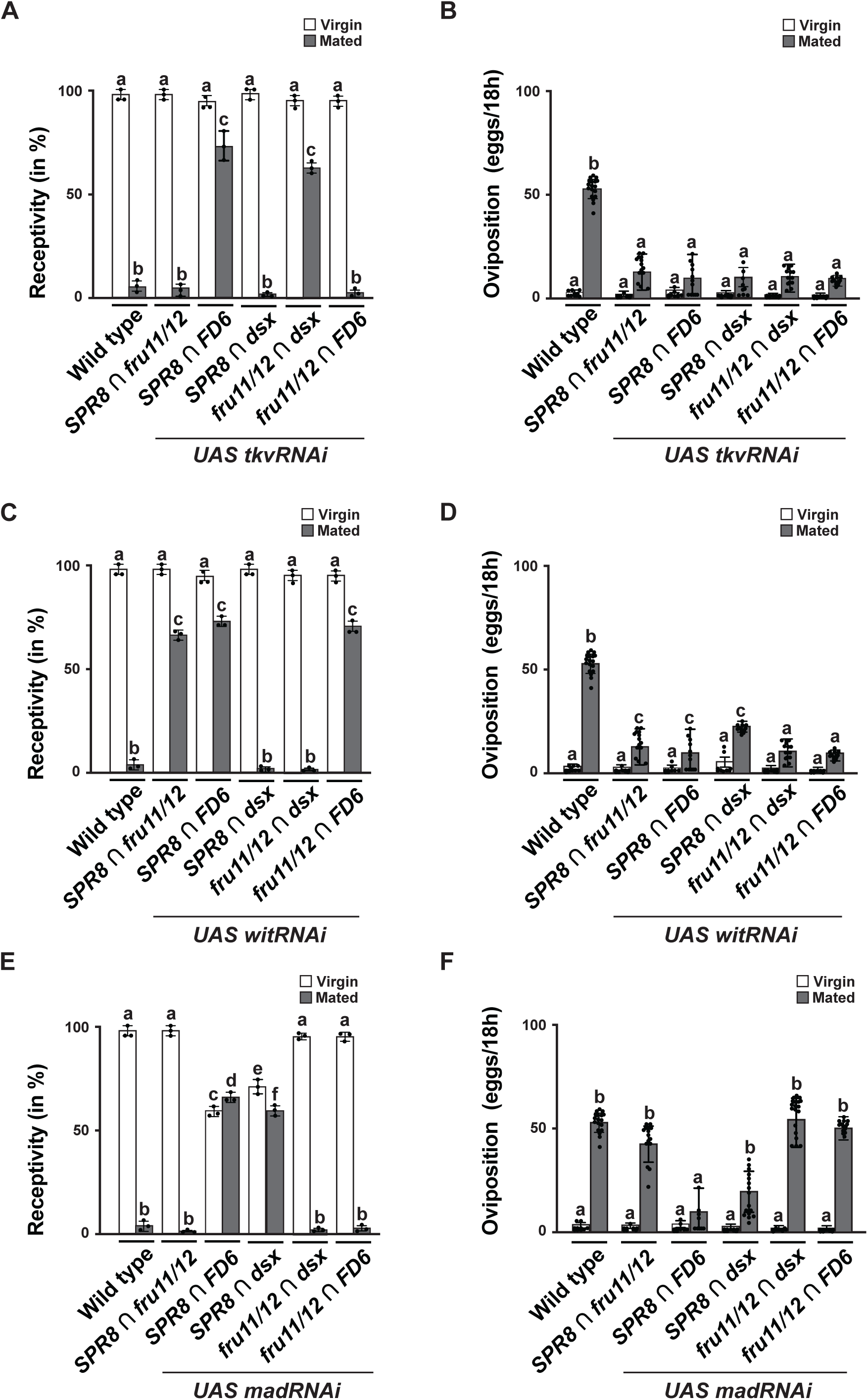
RNAi knockdown of key Dpp/BMP signalling pathway components variably compromises the post-mating response in SPRINz. A-F) Receptivity (A, C and E) and oviposition (B, D and F) of wild type control virgin (white) and mated (black) females expressing *UAS tkvRNAi* (A and B), *UAS witRNAi* (C and D) and *UAS madRNAi* (E and F) under the control of SPRINz *split-GAL4* lines *SPR8 ∩ fru11/12*, *SPR8 ∩ FD6*, *SPR8 ∩ dsx*, *fru11/12 ∩ dsx*, and *fru11/12 ∩ FD6*. Receptivity is shown as mean with standard error from three repeats for receptivity (30 females per repeat) by counting the number of females mating within a 1 h period and oviposition is measured by counting the eggs laid within 18 hours from 20 females (two independent experiments). Statistically significant differences from ANOVA post-hoc pairwise comparison are indicated by different letters (p≤0.0001except p=0.0207 for comparison between e to f in E).

We then turned to the type II receptor *wishful thinking* (*wit*) and its downstream effector *Mad*. *witRNAi* significantly increased remating in *SPR8 ∩ fru11/12*, *SPR8 ∩ FD6* and *fru11/12 ∩ FD6* (Fig. 3C) and, as with *tkv*, reduced oviposition across all five lines (Fig. 3D). With *madRNAi* remating only increased in *SPR8 ∩ FD6* and *SPR8 ∩ dsx* (Fig. 3E), and, unlike *tkv* and *wit*, oviposition was reduced only in the same two lines (Fig. 3F). Overexpression of mad in SPRINz affected receptivity in *SPR8 ∩ FD6* and *SPR8 ∩ dsx,* and induced oviposition in *SPR8 ∩ fru11/12* and *SPR8 ∩ FD6*.

Finally, we downregulated *dpp* itself in the all five SPRINz *split-GAL4* lines. In *SPR8 ∩ FD6*, *fru11/12 ∩ dsx* and *fru11/12 ∩ FD6, dppRNAi* significantly reduced receptivity in virgin females, and also significantly increased remating in *fru11/12 ∩ dsx* and *fru11/12 ∩ FD6* (Supplementary Figure 2A). Oviposition was unaffected in virgin *fru11/12 ∩ dsx* and *fru11/12 ∩ FD6* neurons, but *SPR8 ∩ fru11/12*, *SPR8 ∩ FD6* and *fru11/12 ∩ FD6* females showed significantly reduced oviposition compared to mated controls (Supplementary Figure 2B).

Together, these results show that RNAi of BMP pathway components can disrupt post-mating behaviours in an overlapping, but non-identical subset of SPRINz.

### *dpp* regulates wiring in a subset of SPRINz

Given that *dpp* is a prominent regulator of development, we analysed neuronal wiring of SPRINz in *dpp^HB3^*mutant females using *UAS mCD8GFP* expression. Since *splitGAL4* lines are inserted at *attP40* at 25C, we used this opportunity to remove the lesion in *msl-2* by meiotic recombination (see Methods section) to further ensure that SP insensitivity is not potentiated by the lesions in *msl-2*, which is involved in dosage compensation as part of sex determination (Schutt and Nothiger, 2000), but can also have roles in development (Mineo et al., 2026). The response of these *dpp^HB3rec^*mutant females was indistinguishable from *dpp^HB3^* mutant females upon SP exposure (Fig 4, receptivity: 70±5 %, n=3 and oviposition 3±0.5, n=20)

**Fig. 4:**
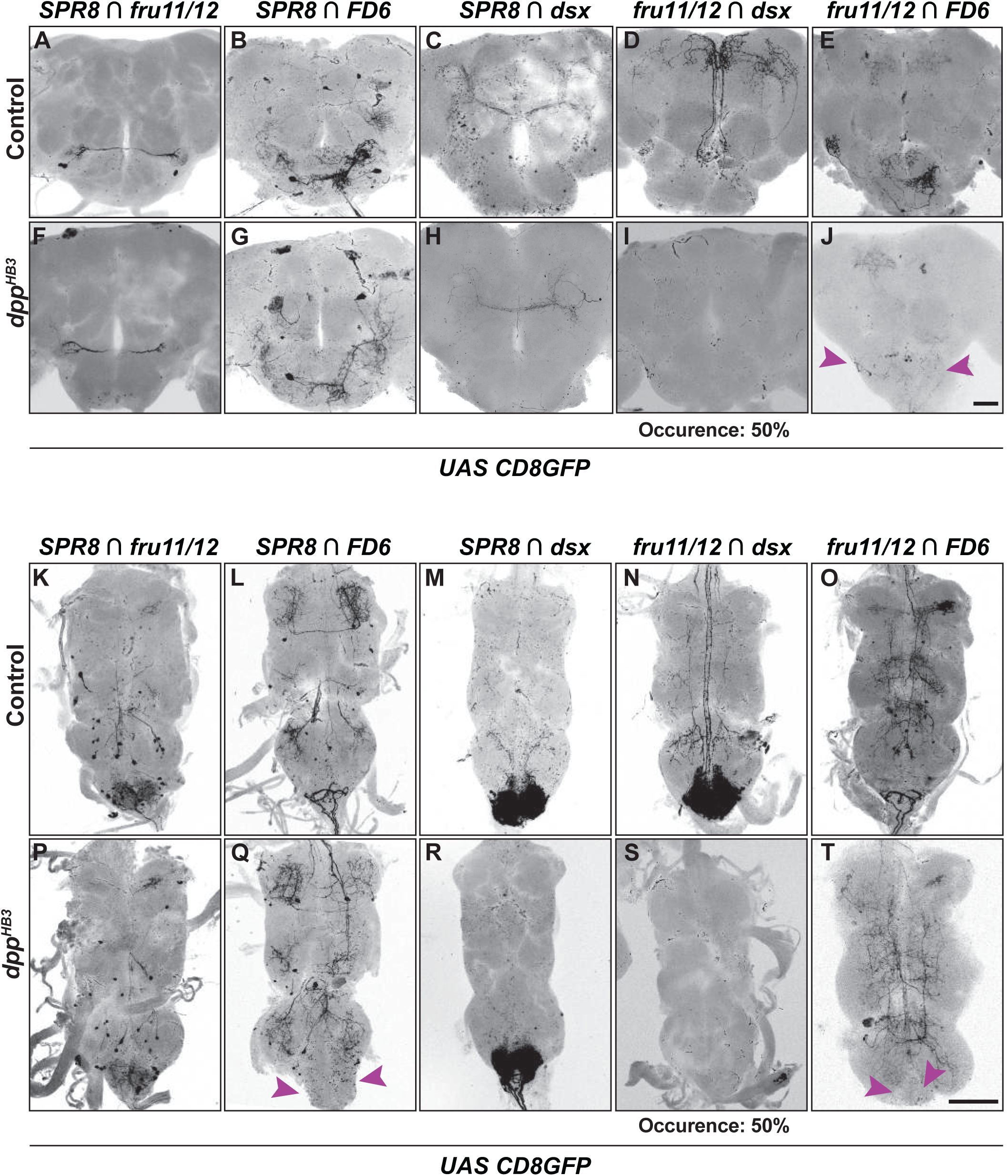
Wiring defects of SPRINz in *dpp^HB3^* females. A-T) Representative adult female brains and ventral nerve cords (VNC) expressing *UAS CD8GFP* under the control of SPRINz *split-GAL4 SPR8 ∩ fru11/12* (brain N=6, VNC N=5)*, SPR8 ∩ FD6* (N=8 brain/vnc)*, SPR8 ∩ dsx* (N=7 brain/vnc)*, fru11/12 ∩ dsx* (N=12 brain/vnc), and *fru11/12 ∩ FD6* (N=9 brain/vnc) in control (*dpp^HB3^*heterozygotes) and *dpp^HB3^* mutants. Note that wiring defects in *fru11/12 ∩ dsx SPRINz* occurred only in 50% of females (n=12, I and S). Scale bars shown in J and T are 50 µm and 100 µm, respectively.

We then compared neuronal projections of SPRINz in *dpp^HB3^*mutant females to control females. Control females displayed stereotypical projection patterns with well-defined axonal trajectories in both CNS and VNC (Fig. 4A-E and 4K-O) (Nallasivan et al., 2026). In contrast, arborizations in 50 % of *fru ∩ dsx* (n=12) and 100 % of *fru ∩ FD6* (n=9) neurons were absent in the *dpp^HB3^* female mutants (Fig. 4I,J). In the abdominal ganglion localized in the ventral nerve cord labelling was absent in *SPR8 ∩ FD6* (n=7), *fru11/12 ∩ dsx* (n=12) and *fru11/12 ∩ FD6* (n=9) SPRINz in *dpp^HB3^* mutant females (Fig. 4S and T).

### Conserved binding motifs for Dpp effector Mad are present in *fru11/12* and *FD6* enhancer fragments

Secreted Dpp binds to BMP receptors that then activate the transcriptional regulator Mad (Wiersdorff et al., 1996; Hudson et al., 1998). Mad binds a GCCGnCGC consensus motif present in multiple Dpp-responsive enhancers (Kim et al., 1997). Some of these enhancers contain motifs present in a tandem arrays to incorporate crosstalk with other signalling pathways (St Johnston et al., 1990).

We therefore examined the regulatory region in the *fru11/12* and *FD6* enhancer to assess whether it contains potential Mad binding sites. Sequence analysis did not detect the canonical GCCGnCGC Mad consensus binding sequences within these regions. However, we find conserved GC-rich tandem motifs in both the *fru11/12* and *FD6* enhancer (Fig. 5A,B).

**Fig. 5:**
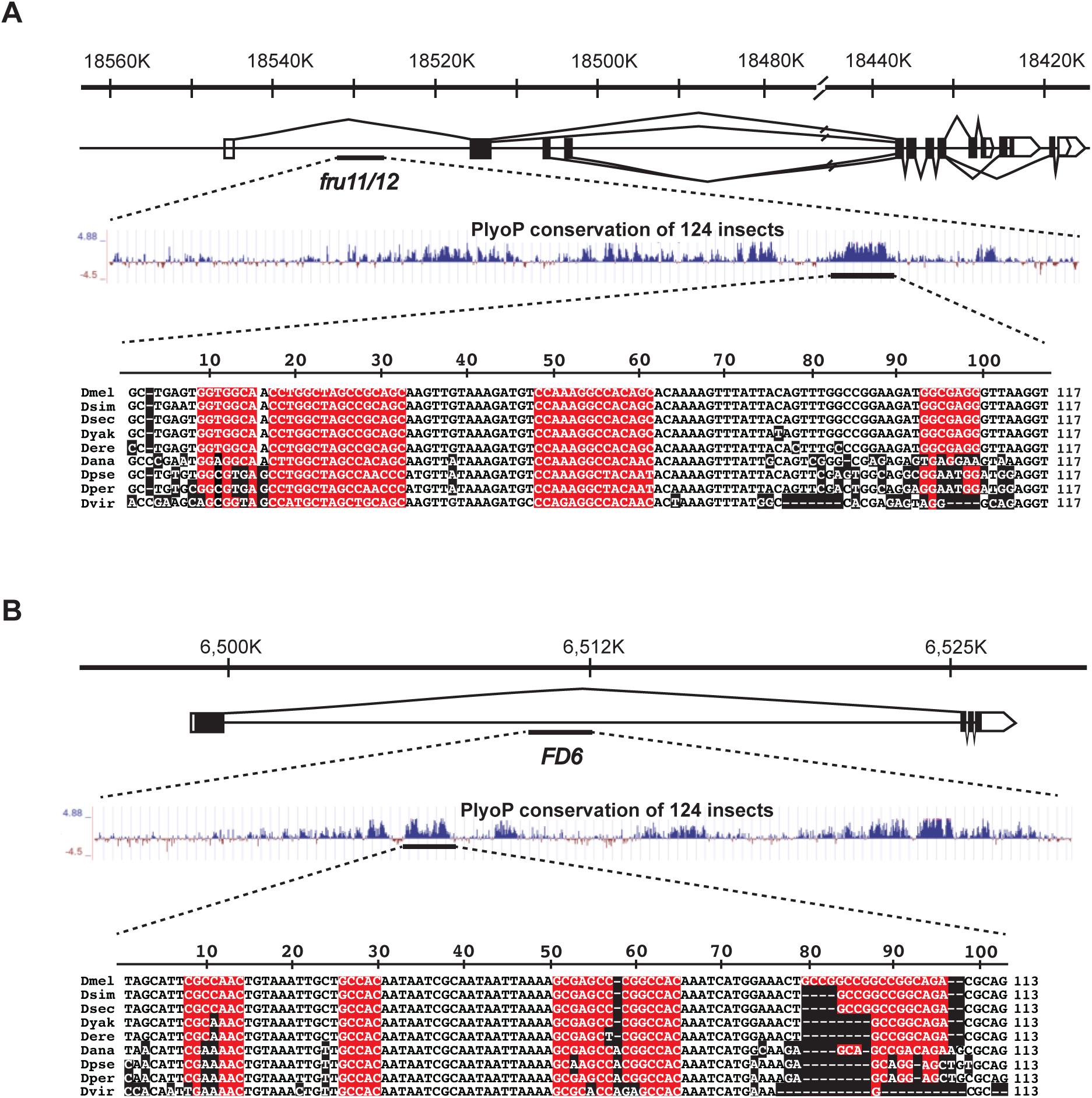
Conserved Dpp response elements are present in the *fru11/12* and *FD6* enhancer fragments. A, B) Sequence alignment of the *fru11/12* and FD6 enhancer fragment in closely related *Drosophila* species (*Dmel*: *D. melanogaster*, *Dsim*: *D. simulans*, *Dsec*:*D. sechellia*, *Dyak*: *D. Yakuba*, *Dere*: *D. erecta*, *Dana*: *D. ananassae, Dpse*: *D. pseudoobscura*, *Dper*: *D. persimilis*, and *Dvir*: *D. virilis*). Nucleic acids deviating from *D. melanogaster* are indicated in black and deletions is dash. The conserved GC-rich motifs of *fru11/12* and *FD6* are indicated in red.

### *Dpp* signalling activates the *fru11/12* enhancer in vivo

To assess whether Dpp signalling activates the *fru11/12* enhancer, we used a dual reporter system consisting of UAS/GAL4 and LexA/LexAOP (Fig. 6A). Here, the *fru11/12 LexA* drives expression in a subset of brain neurons (Nallasivan et al., 2026) that induce tdTomato expression from *LexAop-tdTomato* (Fig. 6A). Then, we expressed Dpp from *UASdpp* pan-neuronally with *elav*-*GAL4* in post-mitotic neurons (Soller and White, 2004). If Dpp can activate transcription through the *fru11/12* enhancer we expect broad neuronal expression of the tdTomato reporter.

**Fig. 6:**
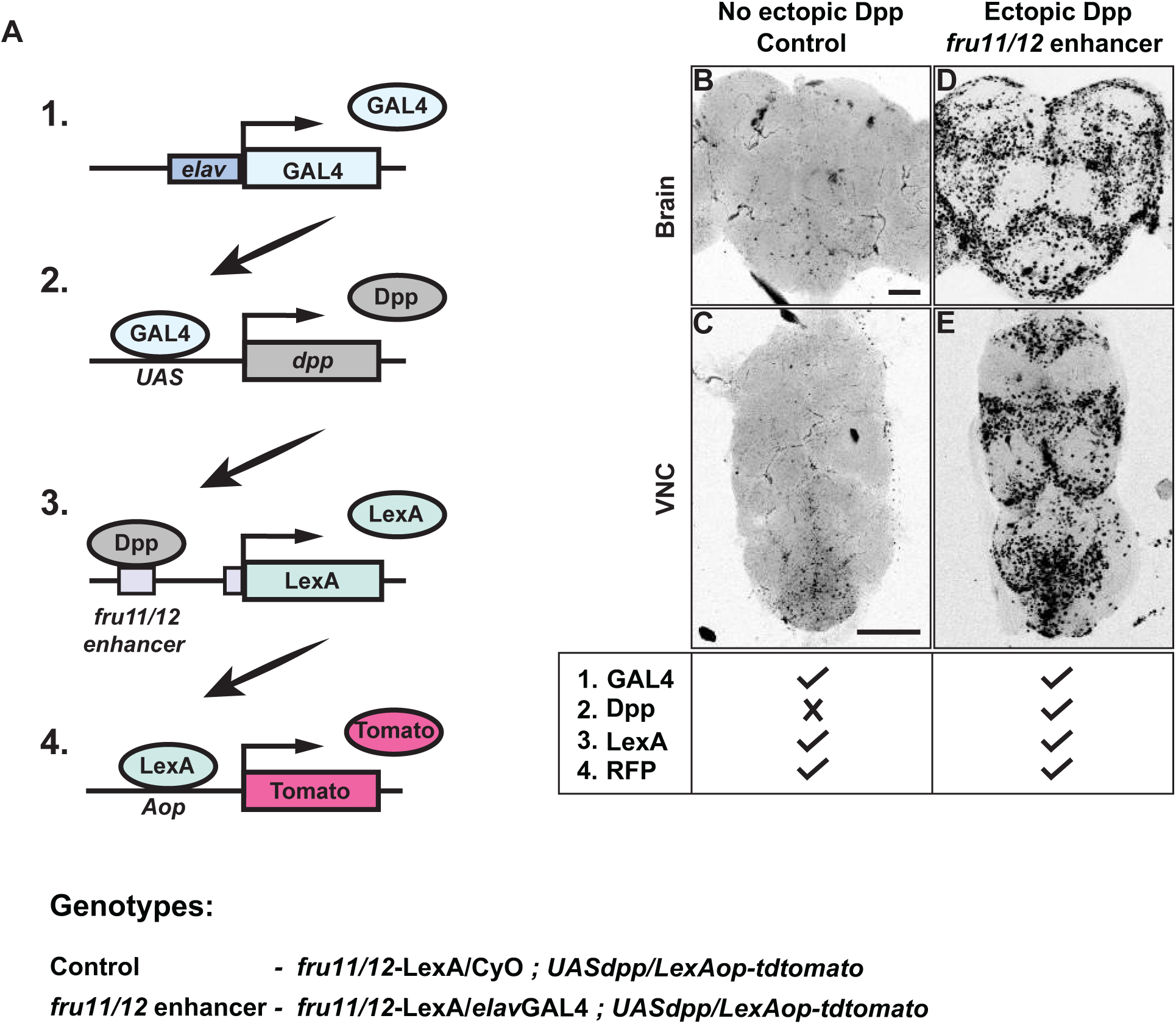
*Dpp* signalling activates the *fru11/12* enhancer in vivo. A) Schematic of the two reporter systems consisting of *GAL4-UAS* and *LexA-LexAop*. The pan-neuronal *elavGAL4* (1) activates transcription from *UAS dpp* (2) to express in post-mitotic neurons. Dpp then binds to the enhancer fragment of *fru11/12* and activates transcription of LexA (3). LexA then induces tdTomato reporter expression from the *LexAop-tdTomato* (4) construct resulting in fluorescent labelling. B-E) Representative adult female brains and ventral nerve cords (VNC) in the absence (B and C) or presence (D and E) of *UAS dpp*. Presence of constructs depicted in A is shown at the bottom. Scale bars shown in B and C are 50 µm and 100 µm, respectively.

**Fig. 7:**
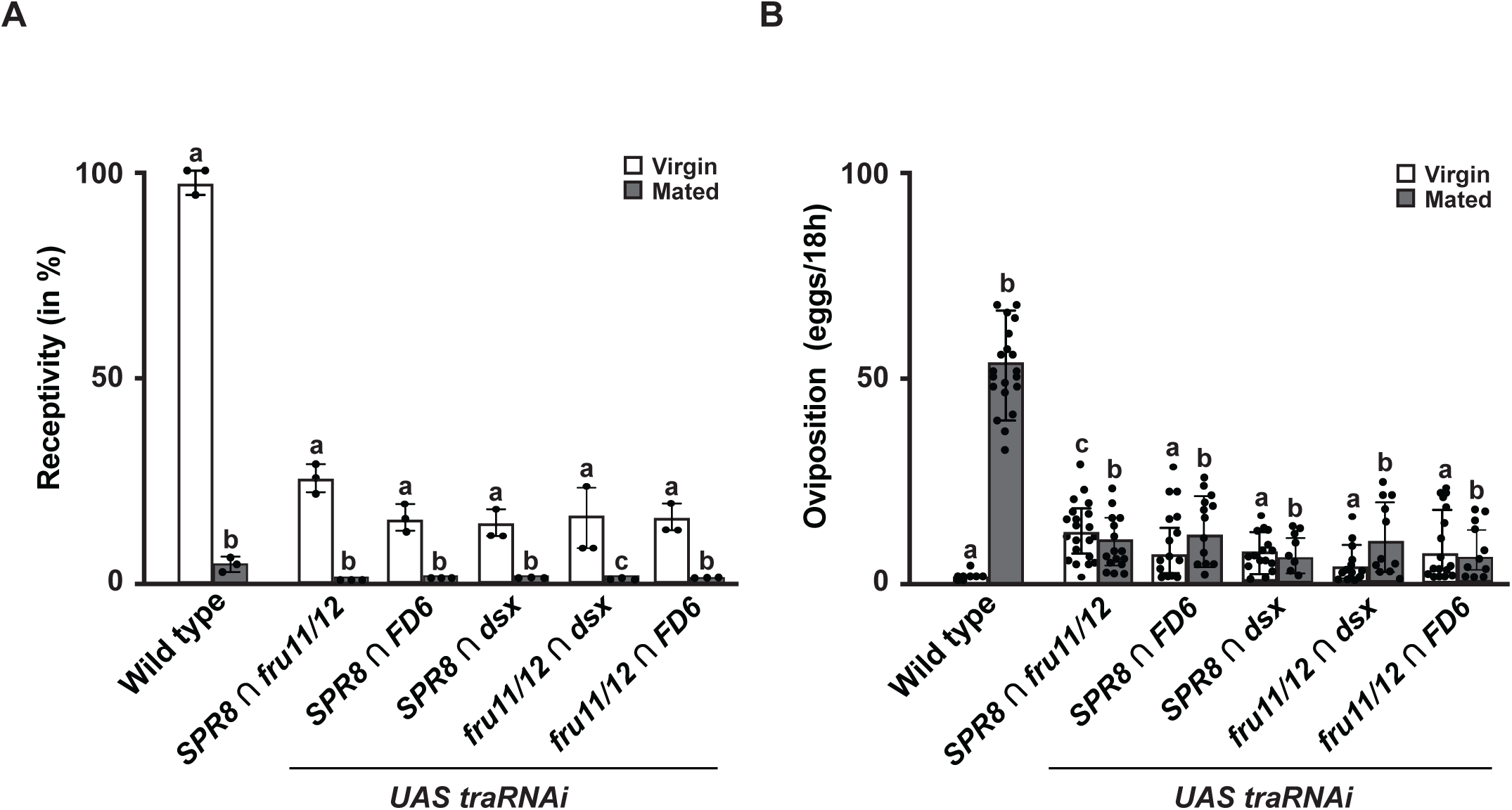
Masculinization of SPRINz disrupts SP induced female post-mating behaviour. A, B) Receptivity (A) and oviposition (B) of wild type control virgin (white) and mated (black) females expressing *UAS traRNAi* under the control of *split-GAL4 SPR8 ∩ fru11/12, SPR8 ∩ FD6, SPR8 ∩ dsx, fru11/12 ∩ dsx,* and *fru11/12 ∩ FD6* lines shown as means with standard error from three repeats for receptivity for virgin females (21 females per repeat) by counting the number of females mating within a 1 h period. For remating of *SPR8 ∩ fru11/12, SPR8 ∩ FD6, SPR8 ∩ dsx, fru11/12 ∩ dsx,* and *fru11/12 ∩ FD6* we obtained 6,4,3,7 and 4 mated females of which none remated. For oviposition by counting the eggs laid within 18 hours from 30 females (two independent experiments). Statistically significant differences from ANOVA post-hoc comparison are indicated by different letters (p≤0.0001).

Indeed, in the absence of ectopiclly expressed Dpp, the tdTomato reporter is only expressed in a few neurons (Fig. 6B AND C). However, if in addition Dpp is express pan-neuronally, we find transcriptional activation through the fru11/12 enhancer broadly in many neurons (Fig. 6D and E).

### Female-differentiation of SPRINz is under the control of the *Drosophila* sex determination pathway

Establishment of sexual identity occurs in a critical window during early pupariation and impacts on the sex peptide response (Arthur et al., 1998). Since defects in *dpp* expression result in neuronal wiring defects in SPRINz neurons we wanted to test whether the *split-GAL4* lines express before SPRINz are differentiated. Therefore, we masculinized SPRINz by removing the female sex determinant Tra by *RNAi*. If the SPRINz *split-GAL4* lines expressed after this critical window, we would not detect changes in the female post-mating response.

*tra RNAi* knock-down in all five populations of SPRINz (*SPR8 ∩ dsx*, *SPR8 ∩ fru*, *SPR8 ∩ FD6*, *fru ∩ dsx* and *fru ∩ FD6*) affected both receptivity and oviposition in virgin females (Nallasivan et al., 2026). Mimicking the SP response, they refused to mate and laid eggs (Fig. 6A,B), indicating that correct sexual differentiation of these neurons is required for the SP induced post-mating response.

## Discussion

Here, we show that the canonical BMP morphogen Dpp plays a critical role in specifying neuronal circuits mediating the SP induced post-mating response. A viable hypomorphic allele of *dpp* (*dpp^HB3^*) displays an egg retainer phenotype and is insensitive to SP. RNAi-mediated knockdown of Dpp/BMP in a subset of SP response inducing neurons (SPRINz) consisting of the *fru11/12* and *FD6* enhancer fragments alters receptivity and oviposition and affects proper wiring in brain and ventral nerve cord neurons. Conserved binding motifs for the Dpp effector Mad are present in the fru11/12 and FD6 enhancer fragments. To validate this, we expressed Dpp ectopically in post-mitotic neurons, which led to broad transcriptional activation of *LexAop-tdTomato* through the fru11/12 enhancer in many neurons, revealing Dpp-induced transcriptional differentiation programs underlying sex-specific behaviours. Likewise, we find that correct sexual differentiation of SPRINz, controlled by the *Drosophila* sex determination pathway, requires Tra for a normal SP-induced post-mating response. Together, these findings establish Dpp and Tra as developmental regulators of neural connectivity underlying female reproductive behaviours.

### The molecular lesion in *dpp^HB3^* in the transcription termination zone affects *dpp* expression

The role of *dpp* in regulating tissue growth, patterning, and cell fate specification is well defined (Entchev et al., 2000; Teleman and Cohen, 2000; Affolter and Basler, 2007; Matsuda et al., 2021; Matsuda and Affolter, 2023). Here, we identified a homozygous viable EMS-induced mutant allele of *dpp* (*dpp^HB3^*) which retains eggs and fails to reduce receptivity following SP exposure (Soller, 2006; Schupbach, 1991). In *dpp^HB3^*, wiring of SPRINz is disrupted adding a new role for *dpp* beyond specifying cell-fate. Similar genetic approaches have previously identified hypomorphic alleles of *egghead* and *Nup54* genes involved glycosphingolipid biosynthesis and nuclear pore function resulting in disruptions of neuronal circuits underlying female post-mating responses (Soller et al., 2006; Nallasivan et al., 2021).

The hypomorphic *dpp^HB3^* allele is homozygous viable, but lethal when one copy is removed, indicating Dpp dosage sensitivity consistent with its role as morphogen. Intriguingly, the molecular lesion in the *dpp^HB3^* allele lies outside the coding region and is at the end of the gene after the polyA site in the transcription termination zone. This lesion affects *dpp* transcript levels in tissue-specific manner. Only few examples of gene expression regulation at the level of Pol II transcription termination have been reported (Soller, 2006; Winczura, 2026). Since 3’end cleavage and polyadenylation precede transcription termination, these two events have been seen as independent with regard to gene expression levels (Kuehner et al., 2011; Proudfoot, 2011). Since there are tRNA genes transcribed by Pol III directly after *dpp*, possible interference could result from these tRNA loci.

### *dpp* regulates neuronal wiring of a subset of SPRINz

Beyond cell fate determination during early development and maintaining germline stem cell identity in the ovary (Affolter and Basler, 2007), Dpp signalling also regulates the development of neuroendocrine structures in the *Drosophila* brain (de Velasco et al., 2007). Dpp secreted along the dorsal midline of the early embryo restricts the formation of the pars intercerebralis (PI) and pars lateralis (PL), two neuroendocrine centres that give rise to neurosecretory cells projecting to the ring gland (de Velasco et al., 2007). Of note, the PI has previously been linked to the SP response by microablation (Bouletreau-Merle, 1976), but direct cellular targets in this brain area have not been identified (Nallasivan et al., 2021).

We observed that RNAi-mediated knockdown of the type I receptor *tkv* in the five SPRINz *split-GAL4* lines increased remating in a subset (*SPR8 ∩ FD6* and *fru11/12 ∩ dsx*), while oviposition was reduced with in all five SPRINz *split-GAL4* lines.

Similarly, knockdown of the type II receptor *wit* increased remating in *SPR8 ∩ fru11/12*, *SPR8 ∩ FD6* and *fru11/12 ∩ FD6* neurons, and likewise reduced oviposition in all five SPRINz *split-GAL4* lines. In contrast, knockdown of the Dpp downstream effector *mad* increased remating with only SPRINz *split-GAL4* lines, *SPR8 ∩ FD6* and *SPR8 ∩ dsx*, where it also reduced receptivity in virgin females and reduced oviposition, while the remaining three populations were unaffected. Finally, knockdown of *dpp* itself reduced receptivity in virgin females and increased remating in *SPR8 ∩ FD6*, *fru11/12 ∩ ds*x and *fru11/12 ∩ FD6* neurons, and reduced oviposition in *SPR8 ∩ fru11/12*, *SPR8 ∩ FD6* and *fru11/12 ∩ FD6* neurons.

Along with Dpp, BMP signalling can also be activated by additional ligands Glass bottom boat (Gbb) and Screw (Scw), which act through distinct heteromeric receptor combinations (Haerry et al., 1998; Neul and Ferguson, 1998; Nguyen et al., 1998). Different tissues and cell types use alternative combinations of receptors and ligands, but crosstalk between ligands can affect Dpp signalling to either potentiate or dampen the outcome (Haerry et al., 1998; Neul and Ferguson, 1998; Nguyen et al., 1998; Rawson et al., 2003). Therefore, the behavioural outcome in response to receptor knockdown is context-dependent, reflecting which ligand and receptor combination a given SPRINz *split-GAL4* relies on. These results also provide further evidence supporting the idea that receptivity and oviposition behaviours are regulated by partially independent neural pathways rather than a single linear circuit (Haussmann et al., 2013; Nallasivan et al., 2021; Nallasivan et al., 2026).

Alongside these behavioural changes we identified wiring defects in *dpp^HB3^* mutants in SPRINz *split-GAL4* labelled neurons. Currently, the tools to resolve synaptic connectivity remains limited but will likely be overcome in the near future by detailed connectome analysis (Galili et al., 2022). Of note, in *D. simulans*, a close relative of *D. melanogaster*, projections of *fruitless P1* sensory neurons that control courtship have changed and alter mate preference (Seeholzer et al., 2018). Likewise, expression of the male Fru isoform in insulin-like peptide producing cells in *Drosophila melanogaster* can trigger nuptial gift giving normally observed in *Drosophila subobscura* (Tanaka et al., 2025). Similarly, wiring defects have also been found in other SP-insensitive mutants of *egghead* and *Nup54* genes (Soller et al., 2006; Nallasivan et al., 2021). Intriguingly, nuclear pore proteins have been associated with speciation (Presgraves, 2010) and the promoter of *Nup54* contains a hot-spot for rapid evolution, possibly through its role in transposon silencing to prevent hybrid dysgenesis (McQuarrie et al., 2023; McQuarrie et al., 2024).

Important in this context, a prominent role for BMP signalling has also been identified in linking neuronal differentiation and synaptic growth at the *Drosophila* neuromuscular junction (NMJ) (de Velasco et al., 2007; Haussmann et al., 2008; Haussmann and Soller, 2010; Sulkowski et al., 2016). BMP signalling promotes synapse homeostasis through both canonical and noncanonical signalling pathways (McCabe et al., 2003; Goold and Davis, 2007; Ball et al., 2010; Kim and Marqués, 2012). Here, canonical BMP signalling retrograde from the muscle to activate presynaptic transcriptional programs that regulate the structural and functional development of synapses at the NMJ (Aberle et al., 2002; Marqués et al., 2002).

Consistent with these findings, we observed wiring defects in SP target neurons in the adult brain and ventral nerve cord (VNC) of *dpp^HB3^* mutants. These defects were more pronounced in *fru ∩ dsx* and *fru ∩ FD6* neurons than in *SPR8 ∩ Fru11/12* and *SPR8 ∩ dsx* populations. In contrast, wiring in *SPR8 ∩ FD6* neurons appeared normal in the brain but displayed defects in the terminal segment of the VNC, specifically within the abdominal neuromere.

Secreted Dpp binds to BMP receptors and activates downstream transcriptional regulators that mediate transcriptional responses (Wiersdorff et al., 1996; Ashe et al., 2000). A key effector of this pathway is Mad, which binds to a conserved consensus sequence GCCGnCGC present in multiple Dpp-responsive enhancers (Kim et al., 1997). Although, we did not identify this exact canonical motif within the enhancer fragments of the *split-GAL4* lines, we detected multiple GC-rich regions within the *fru11/12* and *FD6* enhancer fragments. These GC-rich elements are conserved across *Drosophila* species, indicating functional constraint within these regulatory domains, and in fact we show that Dpp can activate transcription through the *fru11/12* enhancer. Great variability in GC-motifs can be due to differential interpretation of Dpp levels, which results in a graded response as found in a broad analysis of Dpp-responsive enhancers in embryogenesis (Ashe et al., 2000; Wharton et al., 2004; Morikawa et al., 2011).

### Sex-specific differentiation of SPRINz requires Tra

Female sexual differentiation in *Drosophila* is induced by sensing the ratio of X chromosomes to autosomes leading to the expression of the RNA binding protein Sex lethal (Sxl) (Schutt and Nothiger, 2000). Sxl then regulates alternative splicing of *tra* to express Tra only in females for sex-specific alternative splicing of the key transcription factors *dsx* and *fru* leading to male and female differentiation. Differentiation of female neurons occurs during a critical period in early pupariation (Arthur et al., 1998). Since RNAi of *tra* is effective in preventing female differentiation in all SPRINz *split-GAL4* combinations this indicates that these enhancers are turned on before the critical period during pupariation. However, RNAi of Dpp signalling pathway components in SPRINz *split-GAL4* lines affected female sexual differentiation in different ways to respond to SP. Hence it is conceivable, that sexual differentiation is integrated in a broader signalling network required input from other pathways such as Dpp signalling (Clough et al., 2014).

## Materials and Methods

### Fly stocks and genetics

Flies were kept on standard cornmeal-agar food (1%industrial-grade agar, 2.1% dried yeast, 8.6% dextrose, 9.7% cornmeal and 0.25% Nipagin, all in (w/v)) in a 12 h light: 12 h dark cycle. Sexually mature 3–5-day-old virgin females were injected, or mated and examined for their post-mating behaviours as described previously (Soller et al., 1999; Soller et al., 2006). For injections, virgin female flies were cooled to 4° C and 3 pmol SP in 50 nl Ringer’s solution was injected. Ovaries were analysed as previously described (Soller et al., 1999).

The *dpp^HB3^* mutant allele was identified in a genetic EMS screen for oogenesis mutants, whereby one class was characterized with normal oogenesis, but failure of females to lay eggs (Schupbach and Wieschaus, 1991). The genetic background of *dpp^HB3^* was cleaned by recombination with the attP40 phiC31 landing site at 25C containing *fru11* or *SPR8 splitGAL*4 activation domain (AD) inserts. To obtain *dpp^HB3^*attP40 *fru11 or SPR8 AD* recombinants, the attP40 lines were first recombined with either *P{lacW}ND-PDSWk10101* which is a lethal insert next to *msl-2*. In a second step, the lethal insert was then replaced with *dpp^HB3^* by recombination through testing for lethality or sterility with *Df(2L)Exel7011* or *Df(2L)Exel7016*. The *UASdpp* line was a 3rd chromosomal insert *P{UAS-dpp.S}42B.4* (Bloomington #1486). *UAS dppRNAi* lines were *P{TRiP.HMS00011}* (Bloomington #33618), and line *P{TRiP.JF01371}* (Bloomington #25782) was used for dpp RNAi in SPRINz. The UAS *traRNAi* line used was *P{TRiP.JF03132}* (Bloomington #28512) and the SPRINz *split-Gal4* lines were as described (Nallasivan et al., 2026).

### Genome sequencing

For whole genome sequencing of *dpp^HB3^*, high-quality genomic DNA was extracted with the ZR Tissue & Insect DNA MiniPrep kit (Zymo Research) from homozygous *dpp^HB3^* flies as described (Lassota et al., 2025) and 2x150 bp paired-end Illumina sequenced by GATC (Eurofins, Germany) for 5 Mio reads which yields on average 22.5 fold coverage. The demultiplexed data was mapped to the *Drosophila* reference genome (version 6.8) and analysed using IGV by comparison to known polymorphisms from the *Drosophila* Genetic Reference Panel (DGRP2) (Mackay et al., 2012; Huang et al., 2014; Ustaoglu et al., 2021).

For validation genomic DNA was extracted form *dpp^HB3^* as described (Koushika et al., 1999). A distal *dpp* fragment was amplified with primers F1 (GGATCTGGTGAGCAGAGGTTGCGATG) and R1 (GCCGCACAGTTTGGGTTGCCTTTC) and sequenced.

### RNA quantification

Reverse transcription quantitative polymerase chain reaction (RT-qPCR) was done as described by extracting total RNA using Tri-reagent (SIGMA) and RT was done with Superscript II (Invitrogen) according to the manufacturer’s instructions using an oligo dT primer with an anchor sequence and 3’RACE was done as described using primers F1, F2 and AUAP and sequenced with primer F3 (Soller and White, 2005).

For qPCR 1.5 µl cDNA with the SensiFAST SYBR No-Rox kit (Bioline) was used with primers dpp qF1 (CCCCGAGCCGATGAAGAAGCTCTA) and dpp qR1 (GGAATGCTCTTCACGTCGAAGTGCAG), and rpl3F1(CTGGCATCCACACCAAGGACGGCA) and rpl3R1(GGAGCCGATGCAGCAGCCCTTG) (Haussmann et al., 2011). Amplification was done in a Applied Biosystems ABI Prism 7000 with 3 min initial denaturation at 95° C followed by 40 cycles with 15 sec denaturation at 95°C and 60 sec extension at 60° C. Quantification was done according to the ΔCT method as described (Haussmann et al., 2016; Decio et al., 2021).

### Immunohistochemistry and image acquisition

For the analysis of adult axonal projection expressing green fluorescent protein (GFP), tissues were dissected in PBS, fixed in 4% paraformaldehyde in PBS for 15 minutes, washed in PBST (PBS with BSA and 0.3% Triton-x 100).

For imaging, tissues were mounted in Vectashield (Vector Labs) and visualized with confocal microscopy using a Leica TCS SP8. Images were processed using Fiji. Since egg laying and receptivity data are skewed, we used non-parametric Welch’s ANOVA followed by planned pairwise comparisons with Dunnett’s multiple comparison test was done for statistical analysis using Graphpad prism.

## Data availability

All data are available in the main text or the supplementary material as source data in the Figure Source Data file. Reagents are available upon reasonable request from the corresponding author. The *dpp^HB3^* genome sequence has been deposited at GEO (PRJNA1202529).

## Supporting information

Supplemental Figures with figure legends

## Acknowledgments

We thank T. Schüpbach and the Bloomington stock centre for fly lines, D. Scocchia for help with sequencing, D. McQuarrie and R. Arnold for help with sequence analysis and M. Merino for comments on the manuscript. We are indebted to Eric Kubli for his support when this study was initiated. This work was supported by the Biotechnology and Biological Science Research Council to MS.

## Author contributions

MS conceived and directed the project. DNDS performed molecular biology experiments, DNDS and MS performed genetic experiments, DNDS performed imaging and MS performed sequence analysis. All authors analysed data. MS wrote the manuscript with support from DNDS. All authors read and approved the final manuscript.

