## Supplemental Figures with figure legends for "*decepentaplegic* directs wiring of female-differentiated *fruitless* sex peptide response-inducing neurons"

**DEEPANSHU N.D. SINGH AND MATTHIAS SOLLER***

Division of Molecular and Cellular Function, School of Biological Sciences, University of Manchester, Oxford Road, Manchester M13 9PT, United Kingdom

Running title: *dpp* directs neuronal wiring

**Key Words**: *dpp*, *fru*, *tra*, sexual differentiation, neuronal wiring, post-mating behaviors, sex peptide response-inducing neurons (SPRINz)

### Supplementary figure legends

**A**

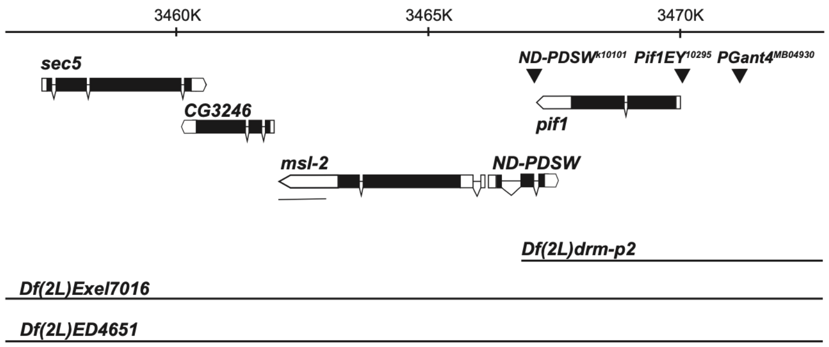


**B**

5'..ATCAACTCCCAAATATCTTTTAGGAGCATTCTACAATCTGATTTTTGTAATCATCCAGCAAACGTTTCCAAAACTCAATTTTTTTTTTAA**A**TACGTATTAAGCCAAAAG

TCGTATACATAACTATTTAAAGCAATTCTGCAGCTGCAGCAAAGATAATCTATTTAAGAGTAAAGCATTAGTCAAAACTAAAGATTATTTTCGTTCAACGAACAGCGAAT

TATAGATTGCTTTGCCCAAAATGTGGATGCTGTGAGCTAAAAAAAACCTTTTTTTTT**AAA**AATAGGCTAAGACATTTTTTTGAATAGGCTAAGACTTTTTTTGA**A**GCAT

GAATTTTTTTTGAGCACGTGAACCTAGGATTAAGAACAACAATTCAAGTAAGCCATATATACCCATATTATATATGCGAACATATAACTATTAAAATATTA**A/C**AATAAAA 3'

### Fig. S1: Multiple nucleotide changes in the Sxl binding sites in *msl-2*

A) Schematic of the *msl-2* chromosomal region depicting gene model and chromosomal deficiencies used below chromosomal nucleotide positions. Transposon insertions are showed as triangles. Coding parts are shown as black and non-coding parts as white boxes. The line under msl-2 indicates the sequence shown in B.

B) Sequence of the msl-2 3'end with changes found in *dpp*$^{HB3}$ in bold and underlined near $U_{\geq 7}$ motif Sxl binding sites and in the polyA site.

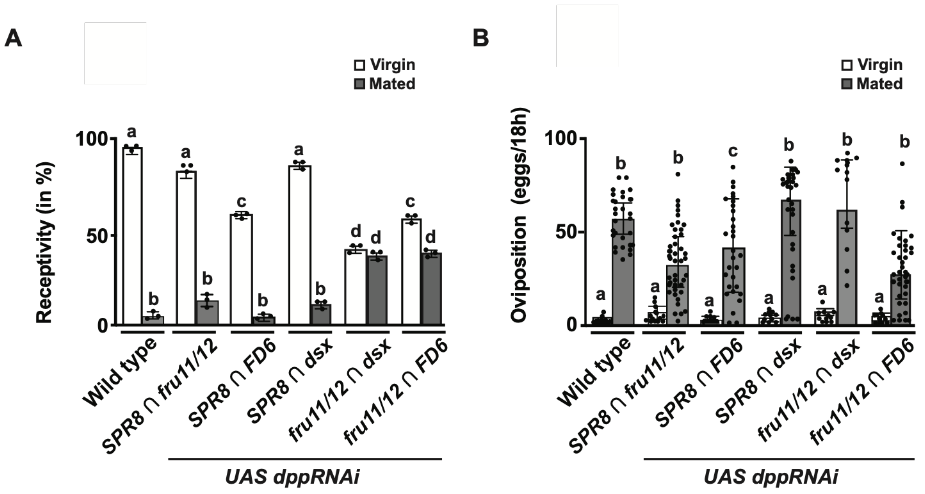


**Fig. S2: *RNAi* knock-down of *dpp* in a subset of SPRINz impacts on the SP response.**

A, B) Receptivity (A) and oviposition (B) of wild type control virgin (white) and mated (black) females expressing *UAS dppRNAi* under the control of *split-GAL4 SPR8 ∩ fru11/12, SPR8 ∩ FD6, SPR8 ∩ dsx, fru11/12 ∩ dsx,* and *fru11/12 ∩ FD6* lines shown as means with standard error from three repeats for receptivity (21 females per repeat) by counting the number of females mating within a 1 h period or for oviposition by counting the eggs laid within 18 hours from 30 females (from two independent experiments). Statistically significant differences from ANOVA post-hoc pairwise comparison are indicated by different letters (p<0.0001 except P=0.003 for c compared to b in B).

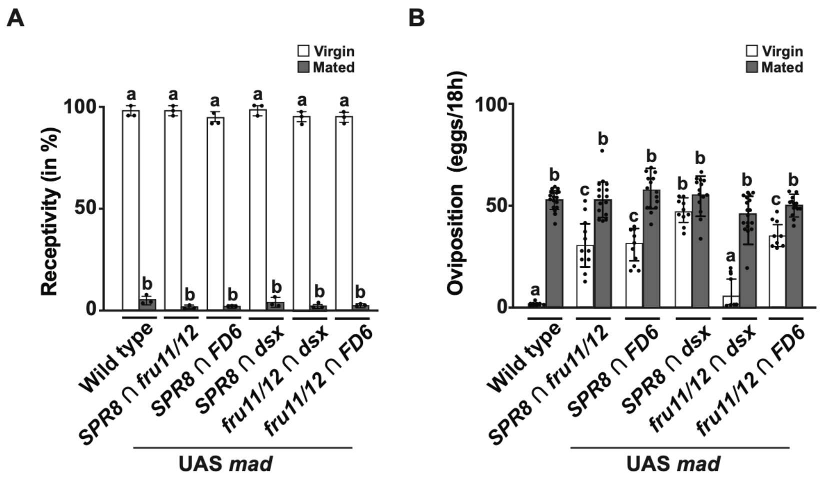


**Fig. S3: Overexpression of *mad* in a subset of SPRINz impacts on the SP response.**

A, B) Receptivity (A) and oviposition (B) of wild type control virgin (white) and mated (black) females expressing *UAS mad* under the control of *split-GAL4 SPR8 ∩ fru11/12, SPR8 ∩ FD6, SPR8 ∩ dsx, fru11/12 ∩ dsx,* and *fru11/12 ∩ FD6* lines shown as means with standard error from three repeats for receptivity (21 females per repeat) by counting the number of females mating within a 1 h period or for oviposition by counting the eggs laid within 18 hours from 30 females (from two independent experiments). Statistically significant differences from ANOVA post-hoc pairwise comparison are indicated by different letters ($p<0.0001$).
